# Determinants and patterns of the endangered brown bear *Ursus arctos* habitat use in the French Pyrenees revealed by occupancy modeling

**DOI:** 10.1101/075895

**Authors:** Blaise Piédallu, Pierre-Yves Quenette, Nicolas Bombillon, Adrienne Gastineau, Christian Miquel, Olivier Gimenez

**Affiliations:** CEFE, CNRS UMR 5175, Université de Montpellier, Université Paul-Valéry Montpellier, EPHE, 1919 Route de Mende, 34293 Montpellier Cedex 5, France; Office National de la Chasse et de la Faune Sauvage, CNERA PAD-Equipe Ours, Impasse de la Chapelle, 31800 Villeneuve-de-Rivière, France; Laboratoire d’Ecologie Alpine, CNRS UMR 5553, Université Joseph Fourier, BP 53, F-38041 Grenoble Cedex 9, France

**Keywords:** dynamic occupancy model, extinction, habitat use, imperfect species detection, large carnivores, local extinction, *Ursus arctos*

## Abstract

The Pyrenean brown bear (Ursus arctos) in the mountainous border between France and Spain is one of the smallest and most endangered populations of large carnivores in Europe. Here, we aimed at assessing trends in brown bear habitat use in the Pyrenees and determining the underlying environmental and anthropogenic drivers. Using detection/non-detection data collected between 2008 and 2014 through non-invasive methods, we developed occupancy models to investigate the dynamic of brown bear habitat use in the Pyrenees accounting for local colonization and extinction processes. First, we found two non-connected occupancy cores, one located in the West and another in the Center of the Pyrenees, with an overall significant decrease in habitat use between 2008 and 2014. Second, we showed a negative correlation between human density and bear occupancy in agreement with previous studies on brown bear habitat suitability. Our results confirm the critically endangered status of the Pyrenean population of brown bears.

## Introduction

Over the last decades, large carnivore populations have been recovering in Europe following the implementation of conservation policies (Chapron *et al.*, 2014). In parallel, conflicts surrounding the animals’ presence subsist (Treves and Karanth, 2003). More than the direct danger caused by carnivore presence, the main sources of conflicts are the damage on livestock and the competition with local hunters (Ericsson and Heberlein, 2003, Gunther *et al.*, 2004, Piédallu *et al.*, 2016a). For these conflicts to be solved or at least mitigated-a necessary step in the conservation of wild populations-the expectations of all stakeholders should be considered and the management decisions should be based on solid ecological data (Redpath *et al.*, 2013).

Among the four species in continental Europe is the brown bear *Ursus arctos*, which is widely distributed all over the continent and split in numerous populations of varying sizes and ranges (Swenson, Taberlet and Bellemain, 2011), including the large Swedish population (Kindberg *et al.*, 2011) or the much smaller one living in the Italian Apennines (Gervasi *et al.*, 2012). One of the smallest and most endangered of these populations resides in the Pyrenees mountains between Southwestern France and Northeastern Spain and is considered to be critically endangered by the IUCN (Huber, 2007). Its survival required the translocation of Slovenian individuals in 1996-97, 2006 and 2016 after only five individuals were detected in 1995, and it remains to this day small and threatened by demographic stochasticity and inbreeding (Chapron *et al.*, 2009, Swenson *et al.*, 2011).

The distribution of a wild population is a key element on which the IUCN relies to determine its conservation status (IUCN, 2012). However, this state variable is difficult to assess in the case of elusive species with large home ranges (Gittleman and Harvey, 1982), brown bear making no exception. To infer the distribution of large carnivores, their monitoring often relies on tracks and indirect observations coupled with DNA analyses to identify the species (e.g., Bellemain *et al.*, 2005, McDonald, 2004, Taberlet *et al.*, 1997). In the case of the French brown bear, its actual distribution remains poorly studied. Martin *et al.* (2012) conducted habitat suitability analyses at a coarse scale on the Cantabrian brown bear population in Spain and applied it in the Pyrenees, and at a local scale using bear detections in the Pyrenees and presence-only methods. Here, we intend to build on these results to address two main issues in standard species distribution models.

First, when dealing with free-ranging populations, species detectability is most likely less than 1, which can lead to false negatives where animals are present but not detected during the survey (Kéry, 2011). Falsely assuming perfect detection can lead to an underestimation of the actual species distribution (Lahoz-Monfort, Guillera-Arroita and Wintle, 2014). Site-occupancy models were specifically developed to explicitly disentangle non-detection from actual absence through the modeling of the imperfect, possibly heterogeneous, observation process (MacKenzie *et al.*, 2002). Second, another limit of standard species distribution models is the assumption that the species always occupy the most favorable area, and that dispersal allows reaching these ideal territories-both statements originating from the ecological niche concept (Leibold, 1995). However, natural barriers or dispersal limitations (such as being an extremely small population) may prevent a species from reaching a favorable area (Araújo and Guisan, 2006). To address this issue, static occupancy models were extended to account for colonization and extinction processes – so-called dynamic or multi-season occupancy models (MacKenzie *et al.*, 2003). Although static occupancy models have often been used on large carnivores (e.g., Bayne, Boutin and Moses, 2008, Carroll and Miquelle, 2006, Carroll *et al.*, 2003, Hines *et al.*, 2010), there are only few applications of dynamic occupancy models (Miller *et al.*, 2013, Molinari-Jobin *et al.*, 2012).

In this study, we identified environmental or anthropogenic drivers and trend in brown bear habitat use in the French Pyrenees. To do so, we fitted dynamic occupancy models to detection/non-detection data obtained through a multi-source systematic monitoring protocol between 2008 and 2014.

## Material & Methods

### 1. Study area and bear population

This study was performed on the French side of the Pyrenees at the border between Northeastern Spain and Southwestern France (Figure 1). The bears that live here mostly descend from individuals that were translocated from Slovenia to the Pyrenees in 1996-1997 (2 females and 1 male) and 2006 (4 females and 1 male), even though one bear’s mother belonged to the remnant of the original Pyrenean bear population which was thought to include 5 individuals in 1995. Field observations suggest that two population cores exist on the French side of the Pyrenees: two male bears have been accounted for in the Western core, and the Central one accounts for the rest of the population (Figure 1).

**Figure 1:**
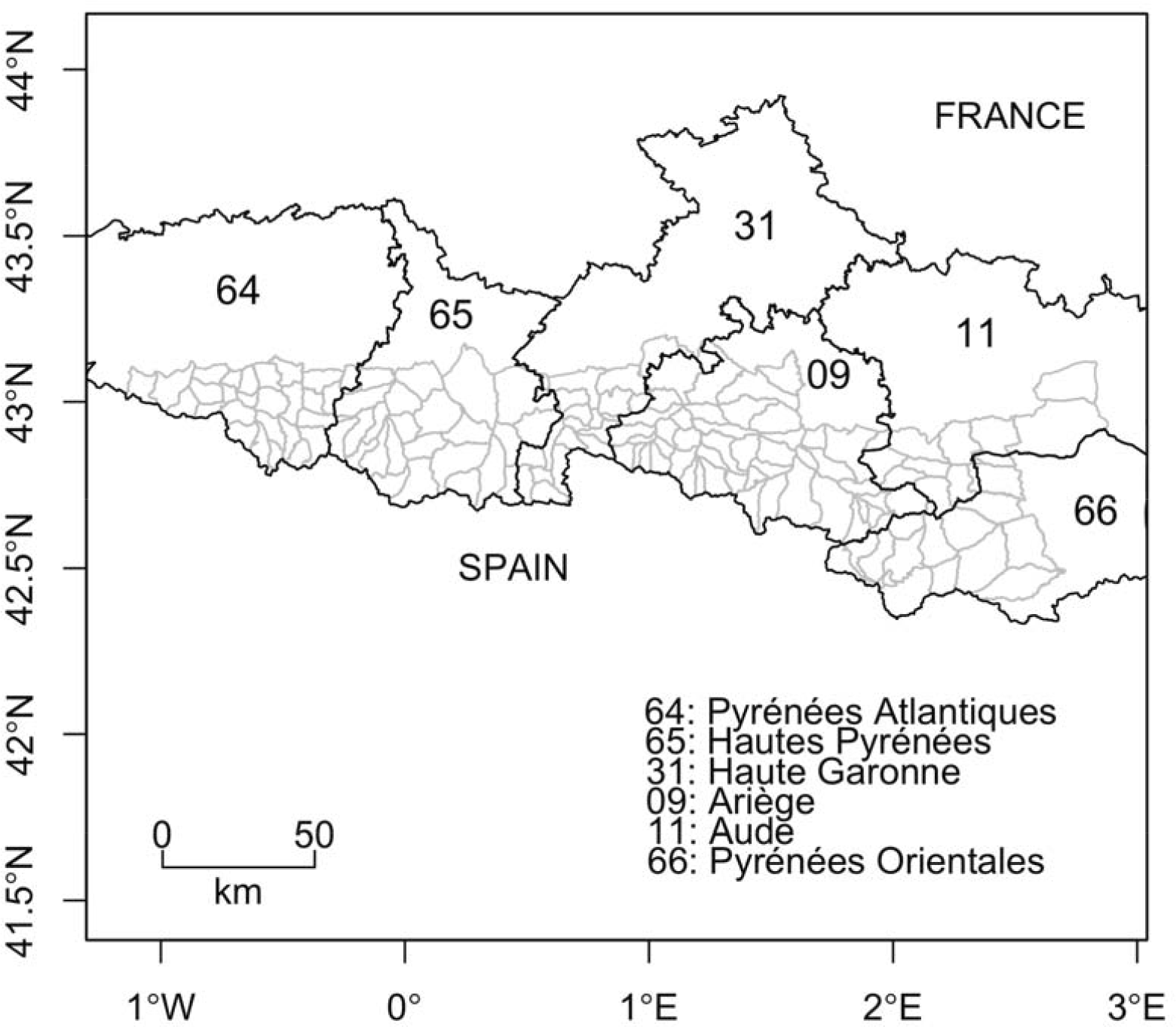
Map of the counties and mountain subsections in the French Pyrenees. Dark lines: county borders. Gray lines: limits between mountain subsections.

### 2. Bear data collection and monitoring

The data used for this analysis was gathered between 2008 and 2014 by members of the national Brown Bear Network (135 professional members from government agencies and 228 unaffiliated amateur members) under the supervision of the French Game and Wildlife Agency (ONCFS). A systematic monitoring protocol was followed using fixed itineraries along which the agents looked for bear tracks such as hair, scats, claw marks or paw prints. The Pyrenees were broken down in mountain massif subsections using ridgelines and the bottom of valleys. The area of a subsection was 95km^2^ on average. This is coherent with the home range of brown bear males and females that is approximately 85-200km^2^ and 50-100km^2^ respectively (Huber and Roth, 1993, Preatoni *et al.*, 2005). Besides, the upper limits of these home ranges were obtained over a period including the rut in May-June, while we used in our analyses the July-November period (see next section) during which the size of home ranges are much reduced because the rut is excluded (Preatoni *et al.*, 2005). Each one of the 84 investigated subsections of the mountain massif included one itinerary. Each itinerary was visited at least once every month from July to November. The length of each itinerary was proportional to the area of the corresponding subsection so as to survey 0.2 km per km^2^ of subsection. Tracks and observations were validated by ONCFS experts, therefore minimizing the risk of false positives due to species misidentification (Molinari-Jobin *et al.*, 2012).

### 3. Model building and selection

To estimate the probability of bear presence in all the mountain massif subsections, we built a dynamic occupancy model (MacKenzie *et al.*, 2003) that was parameterized with the probabilities of colonization γ (the probability for a subsection to become occupied while it was unoccupied the year before), extinction ε (the probability for a subsection to become unoccupied while it was occupied the year before) and initial occupancy ψ_1_ (the probability for a subsection to be occupied the first year of the study), along with the species detection probability p (the probability to detect the species on a subsection when present). We used years as primary occasions, between which colonization and extinction probabilities could be estimated, and the months of July to November as secondary occasions during which we considered the subsections’ occupancy status to remain unchanged (the closure assumption). By focusing on the July-November period, we excluded the reproduction season (April to June) during which male bears in particular are known to increase their movement range while they look for females (Clevenger, Purroy and Pelton, 1990). Despite this precaution, movements may still occur in and out the subsections and, assuming these movements are random, occupancy should be interpreted as habitat use rather than the proportion of area occupied by the species (MacKenzie and Nichols, 2004). More precisely, “the usage made of various habitat components within the home range” is usually referred to as third-order selection (Johnson, 1980).

We relied on previous habitat suitability studies on brown bears in Europe to select candidate environmental and anthropogenic covariates for our analysis (Martin *et al.*, 2010, Martin *et al.*, 2012, Mertzanis *et al.*, 2008). We considered five environmental and anthropogenic covariates for each mountain massif subsection (Table 1; Figure A1). Roughness was obtained as the mean of the absolute differences between the altitude of a massif subsection and the value of its contiguous mountain subsections (Wilson *et al.*, 2007). We used the IGN BD_ALTI^®^ database (250m resolution) to calculate the mean altitude of each massif subsection. Forest cover and shrub cover covariates were extracted from the CORINE Land Cover^®^ database (U.E – SoeS. Corine Land Cover 2012). Road length was built using the IGN ROUTE 500^®^ database. Human density was obtained from the NASA Socioeconomic data and applications center (http://sedac.ciesin.columbia.edu/data/set/gpw-v3-population-count/data-download). The maximum correlation between these covariates was 0.51 in absolute value. We used the Akaike’s Information Criterion (AIC, Burnham and Anderson, 2002) for covariate selection. To compare model support with reference to the model best supported by the data, we used the difference in AIC (ΔAIC). To account for model selection uncertainty, we resorted to model averaging considering all models with ΔAIC < 2. Due to the large number of covariate combinations, we used a multi-stage approach to model selection (Dugger, Anthony and Andrews, 2011, Lee and Bond, 2015, MacKenzie *et al.*, 2012), which proceeded as follows. First, we started by selecting the best model structure by focusing on time-varying covariates only, namely *year* and *survey*. We considered 8 different models in total, with either no effect (.) or a *year* effect on colonization γ and extinction ε, and either no effect (.) or a *survey* effect (where a survey refers to a month, hence a survey specific covariate) on detection probability p (Table 2). Because the sampling effort was homogeneous over the study period, we did not consider a *year* effect on detection. Second, based on previous bear occupancy studies (Martin *et al.*, 2010, Martin *et al.*, 2012, Mertzanis *et al.*, 2011, Nielsen *et al.*, 2010, Nielsen, Stenhouse and Boyce, 2006) and bear biology, we considered *specific* combinations of the environmental or anthropogenic effects on each of the parameters (ψ_1_, γ, ε and p, Table 1). We tested possible negative effects of covariates human density and road length on initial occupancy ψ_1_ as a previous study showed that bears avoided human-caused disturbances (Martin *et al.*, 2010, Mertzanis *et al.*, 2011, Naves *et al.*, 2003). Roughness, shrub cover and forest cover were all positively associated with bear presence albeit performed at different scales in previous studies (Apps *et al.*, 2004, Martin *et al.*, 2010, Martin *et al.*, 2012, Naves *et al.*, 2003, Nellemann *et al.*, 2007). For colonization γ, we studied possible effects of roughness and human density that were the most commonly significant covariates in previous bear distribution studies (Martin *et al.*, 2010). We considered for extinction ε the possible effect of the two anthropogenic covariates human density and road length. Finally, we tested the possible effect of human density, roughness and forest cover on detection p as both could potentially influence the accessibility of bear tracks to observers. To account for the variability in the size of a subsection, we also included its area as a covariate on detection in all models without submitting it to selection. In total, we fitted all possible 8192 models.

**Table 1:**
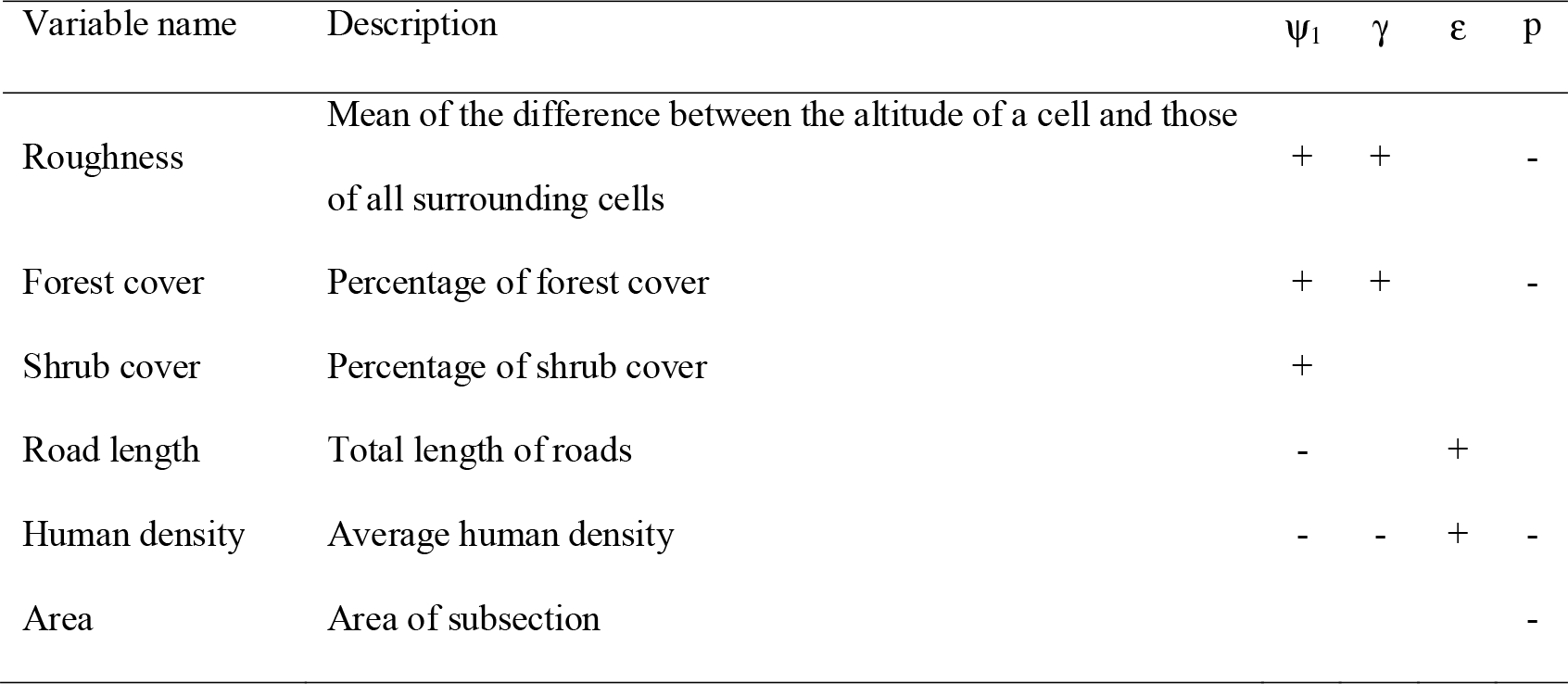
Definition of the environmental variables used for the occupancy analysis, and the parameters for which an effect was tested. ψ_1_: initial occupancy probability, γ: colonization probability, ε: extinction probability, p: detection probability. +/−: predicted sign of the effect of the covariate on the parameter based on previous studies (see text for references). An absence of a +/− sign means that the effect was not tested.

**Table 2:**
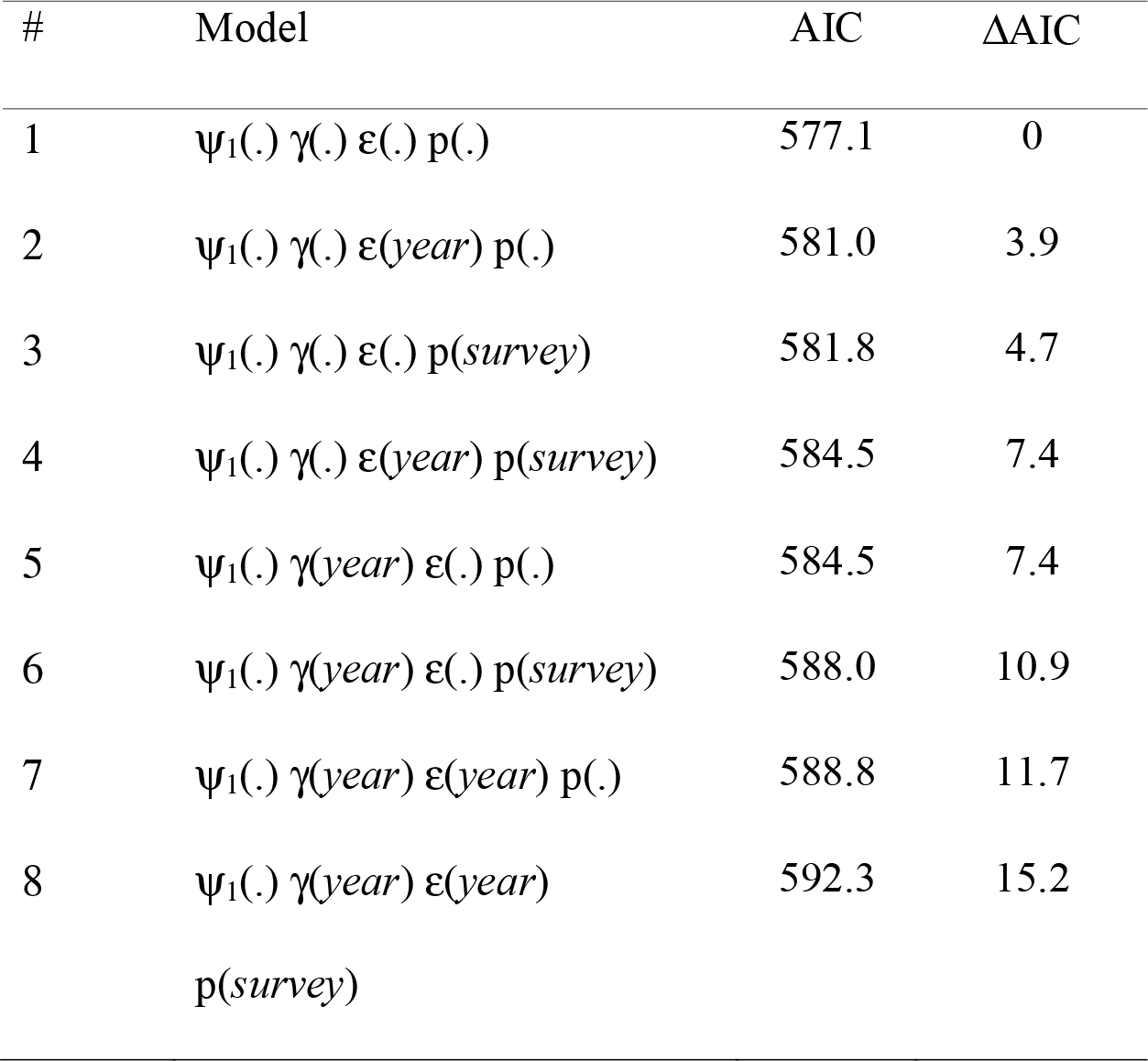
Model selection with time-varying covariates. Models were ranked with AIC. ψ_1_: initial occupancy probability, γ: colonization probability, ε: extinction probability, p: detection probability. *year*: year effect on the parameter, which relates to changes between primary occasions, i.e. from one year to another in our case. *survey*: survey effect on the parameter, which relates to the secondary occasions repeated within a year. ΔAIC: difference between the AIC of the current model and the AIC of the model with lowest AIC.

Because there was uncertainty in the selection of the best set of covariates, we resorted to model-averaging for parameter estimation and inference (Burnham and Anderson, 2002). Effect sizes were examined to determine the magnitude of a covariate effect (Nakagawa and Cuthill, 2007). The covariates were standardized prior to the analyses.

To assess a potential trend over the years in habitat use, we first estimated the occupancy status of each subsection for each year. We then tested a linear effect of year on the binary occupancy variable using a conditional autoregressive correlation model and an adjacency matrix between the different subsections to specify the correlation matrix (Rousset and Ferdy, 2014). A likelihood ratio test (LRT) was performed to assess the significance of this temporal trend. We applied this procedure to all models with ΔAIC < 2.

Eventually, we built maps for initial occupancy, detection, colonization and extinction by calculating the probability at a given subsection using the model-averaged parameter estimates and the value of the covariates for this given subsection.

These analyses were performed in R (RCoreTeam, 2013) with the ‘unmarked’ (Fiske and Chandler, 2011), spaMM (Rousset and Ferdy, 2014), rgdal, AICcmodavg, classInt, RColorBrewer and spdep packages. The data and R codes are available on GitHub at https://github.com/oliviergimenez/ursus_Pyrenees_occupancy.

## Results

### 1. Multi-stage model selection

We found no *year* or *survey* effects on any of the parameters ψ_1_, γ, ε or p (Table 2). The ΔAIC of the next two best models (with a *year* effect on extinction ε and a *survey* effect on detection p respectively) was >2, therefore we used the model with constant parameters as the basic structure for the next step. Despite model uncertainty in the results of the selection procedure on environmental and anthropogenic covariates, some covariates were always included in models with ΔAIC < 2 (Table 3): roughness on detection and colonization probabilities and human density on extinction and initial occupancy probabilities.

**Table 3:**
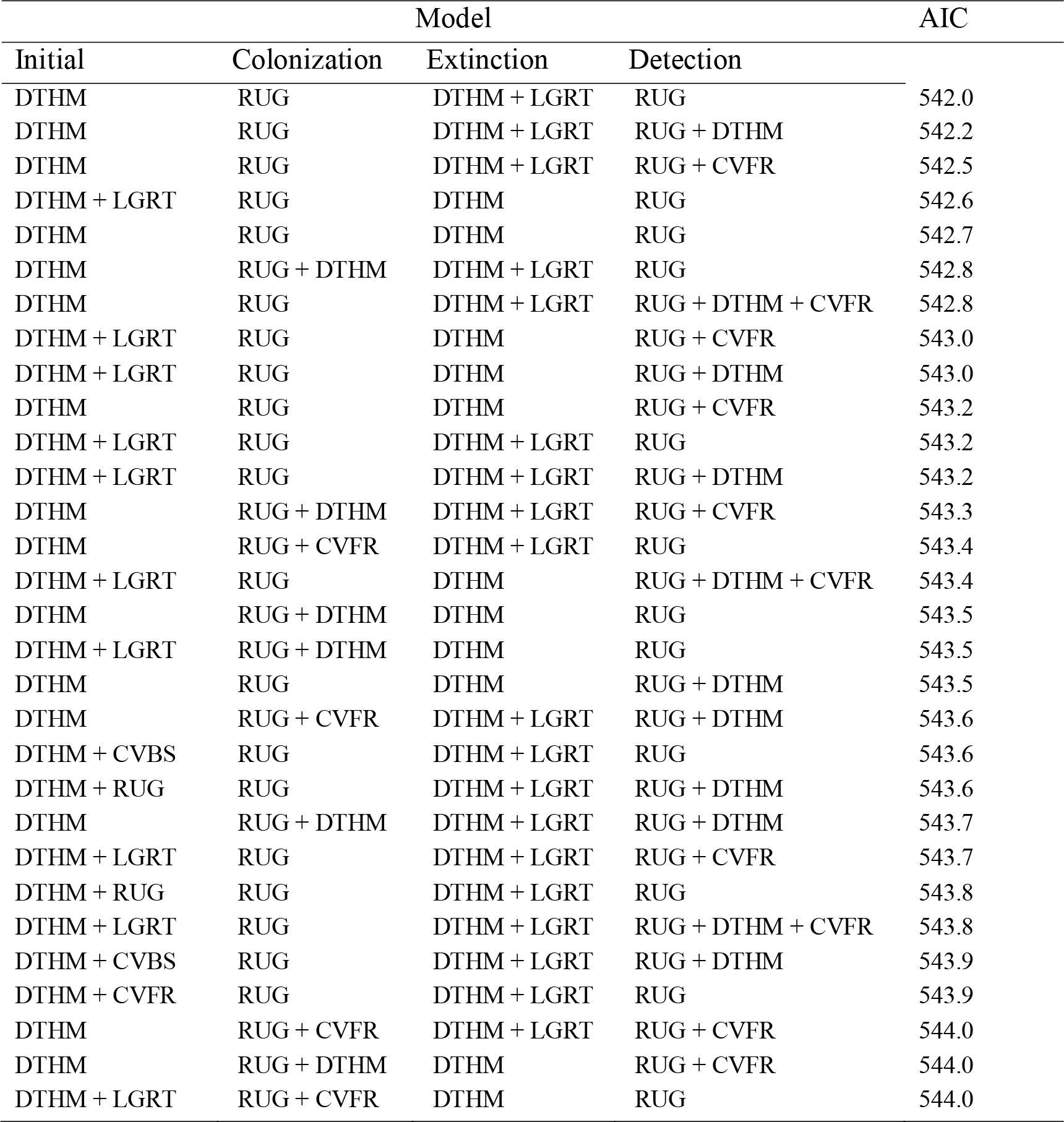
Model selection with environmental and anthropogenic covariates. The covariates and their combinations are detailed in Table 1. Among the 8192 models that we fitted, we only report models with ΔAIC < 2. DTHM is for human density, RUG is for roughness, CVFR is for forest cover, LGRT is for road length, CVBS is for shrub cover. Note that the area of subsections was used in all models in the detection probability and not subject to the covariate selection procedure.

### 2. The effect of covariates on parameters

We refined the patterns found in the covariate selection step by examining the effect sizes (on the logit scale; Figure 2). While the effect of roughness on detection probability and that of human density on both extinction and initial occupancy probabilities were confirmed, the colonization probability was not associated with any covariates. We investigated further the links between the covariates and initial occupancy, colonization, extinction and detection probabilities by assessing the shape of these relationships (after back-transformation; Figure 3). An increase in roughness was associated with an increase in the detection probability, while it was more difficult to detect bears (when present) in large subsections. Initial occupancy ψ_1_ was strongly negatively correlated with human density (Figure 2B), with the least populated areas being much more likely to be used by bears, just like extinction ε was negatively correlated with human density.

**Figure 2:**
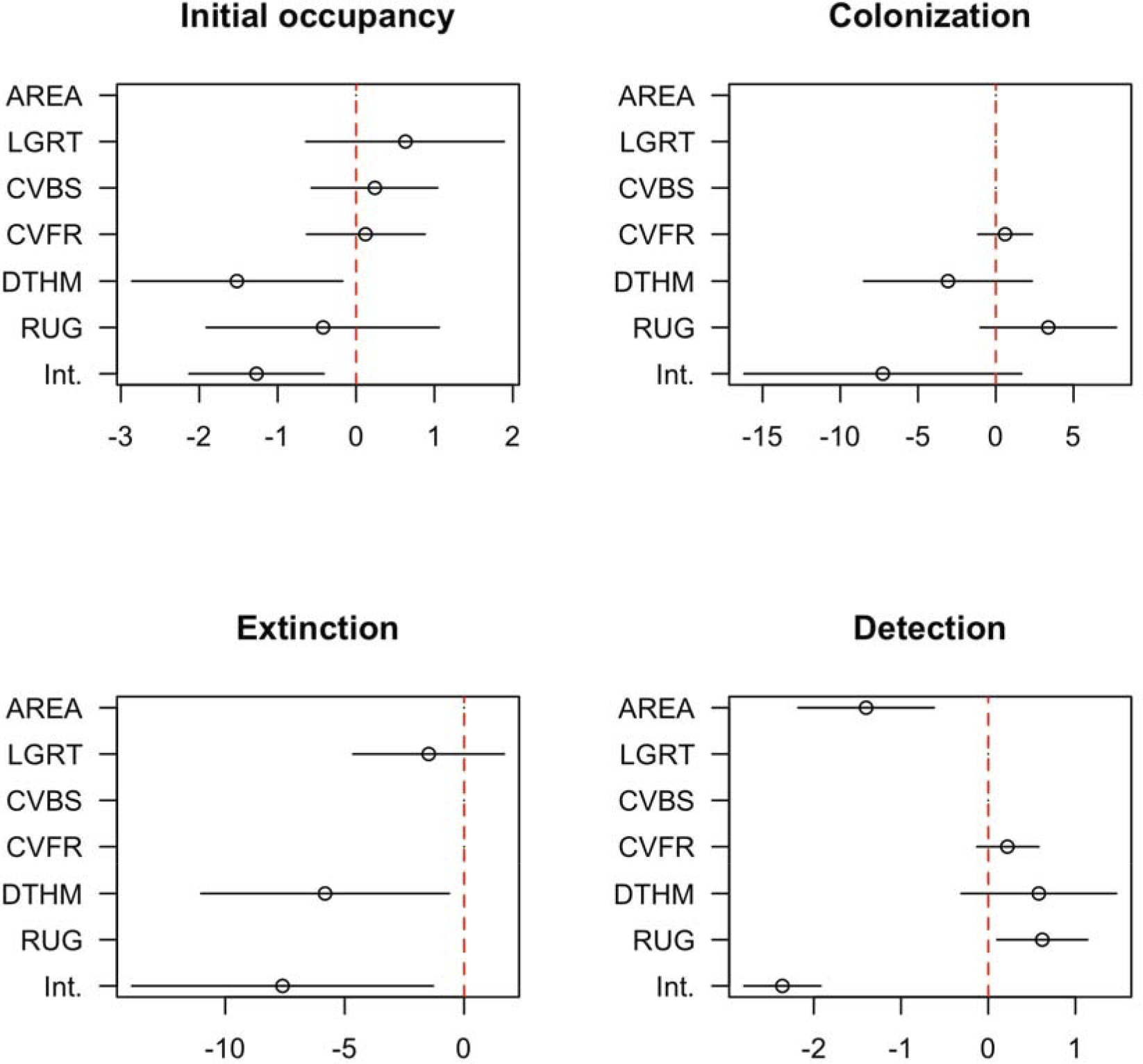
Model-averaged parameter estimates (on the logit scale) and confidence intervals of the (standardized) covariates effects on initial occupancy, colonization, extinction and detection probabilities. An effect does not appear if the corresponding covariate was not considered in the selection procedure. The covariates and their combinations are detailed in Table 1: Int. is for the intercept, DTHM is for human density, RUG is for roughness, CVFR is for forest cover, LGRT is for road length, CVBS is for shrub cover and AREA is for the area of subsections. Note that AREA was used in all models in the detection probability and not subject to the covariate selection procedure.

**Figure 3:**
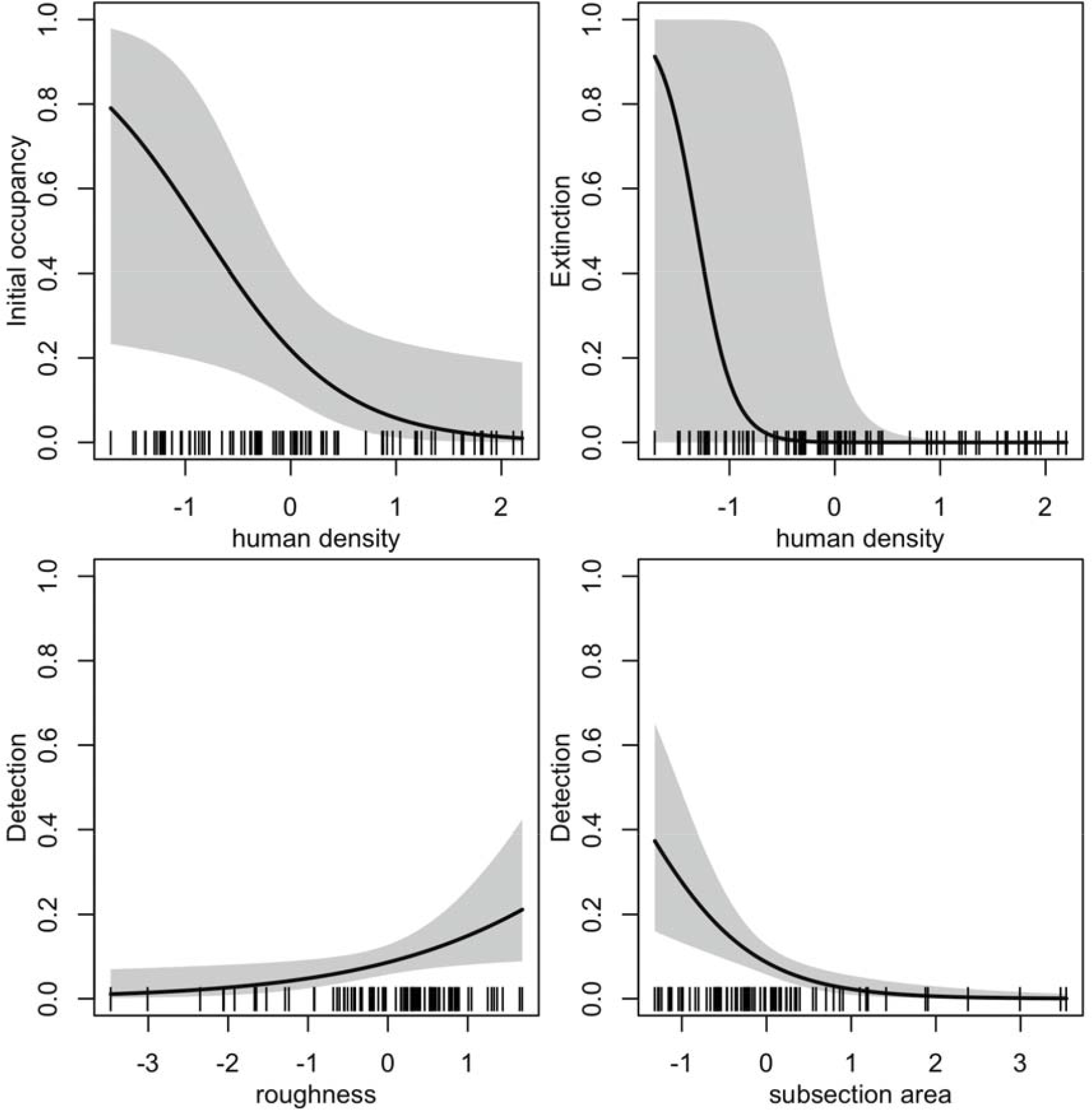
Relationships between model-averaged parameter estimates and standardized covariates. We focused on the most important covariates based on the effect sizes given in Figure 2. The parameter of interest is on the y-axis, namely initial occupancy, extinction and detection probabilities. The colonization probability is not displayed because of small effect sizes (we refer to Figure A2 for the relationships between parameters and all covariates). For each relationship, the non-focal covariates were set at their mean. The dashes on the x-axis indicate the observed covariate values.

### 3. Distribution maps

The initial occupancy map (Figure 4B) clearly showed two population cores (Western and Central), with the Central Core extending in Southeast Ariège and Southwest Aude and Pyrénées-Orientales (Figure 1). The extinction probability in the East of the Central core was high (Figure 4D), which is consistent with the disappearance of the bears from that area (Camarra *et al.*, 2012), while the colonization probability in the same mountain subsections were close to zero (Figure 4C). Detection was higher in the Central core than it was in the Western core (Figure 4A). The colonization map indicated that the Western population core was more likely to expand to the East, while the Central one was more likely to expand to the West (Figure 4C). These last observations were confirmed by the yearly occupancy maps (Figure 5), which showed a decrease of the occupancy probability in the Eastern parts of the Central population core (Southeast Ariège, Southwest Aude and Pyrénées-Orientales).

**Figure 4:**
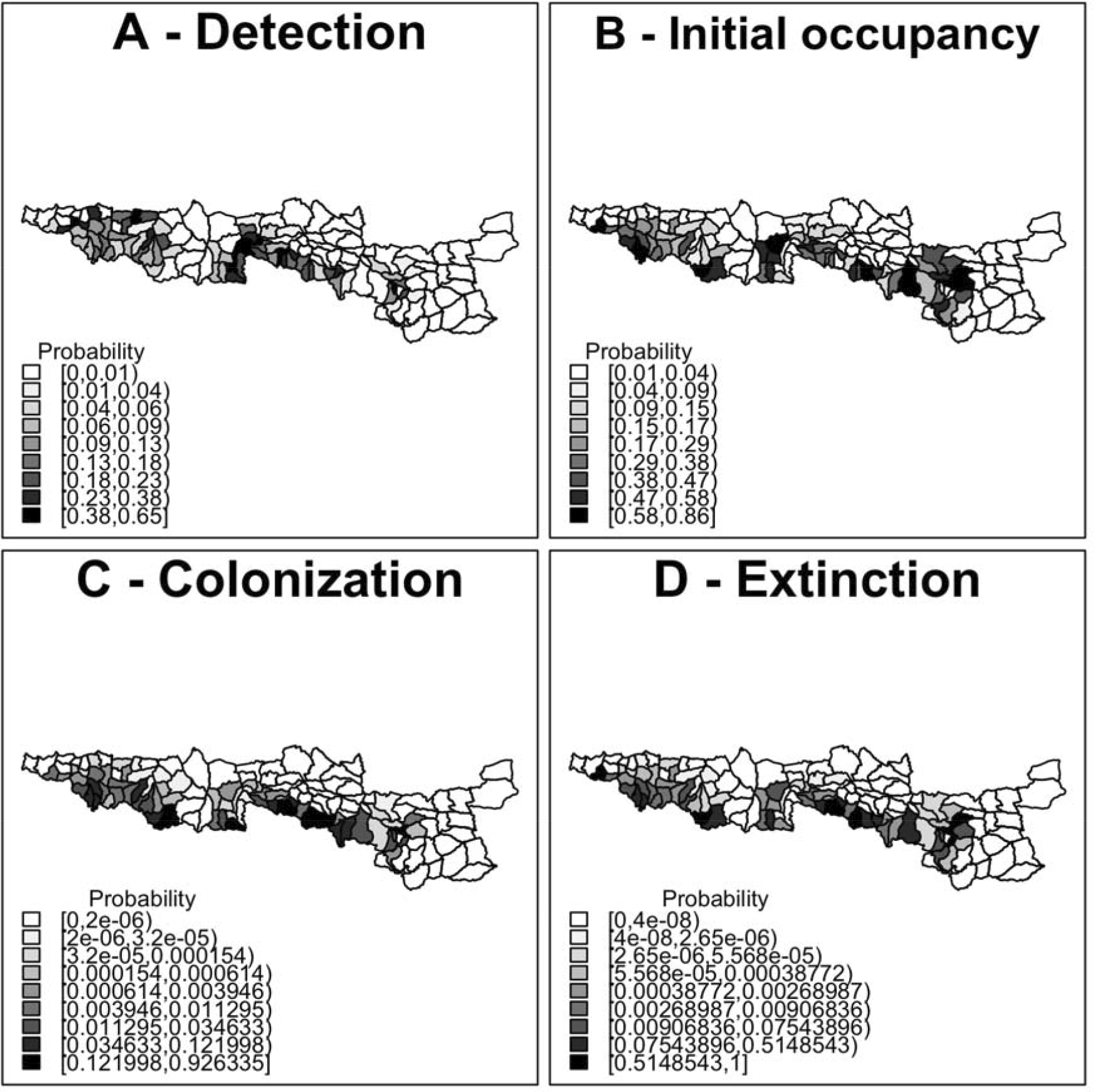
Maps of the model parameters in the various mountain subsections of the French Pyrenees obtained using the model-averaged parameter estimates. A: Detection probability, B: Initial occupancy probability, C: Colonization probability, D: Extinction probability. Covariates were set at their mean.

**Figure 5:**
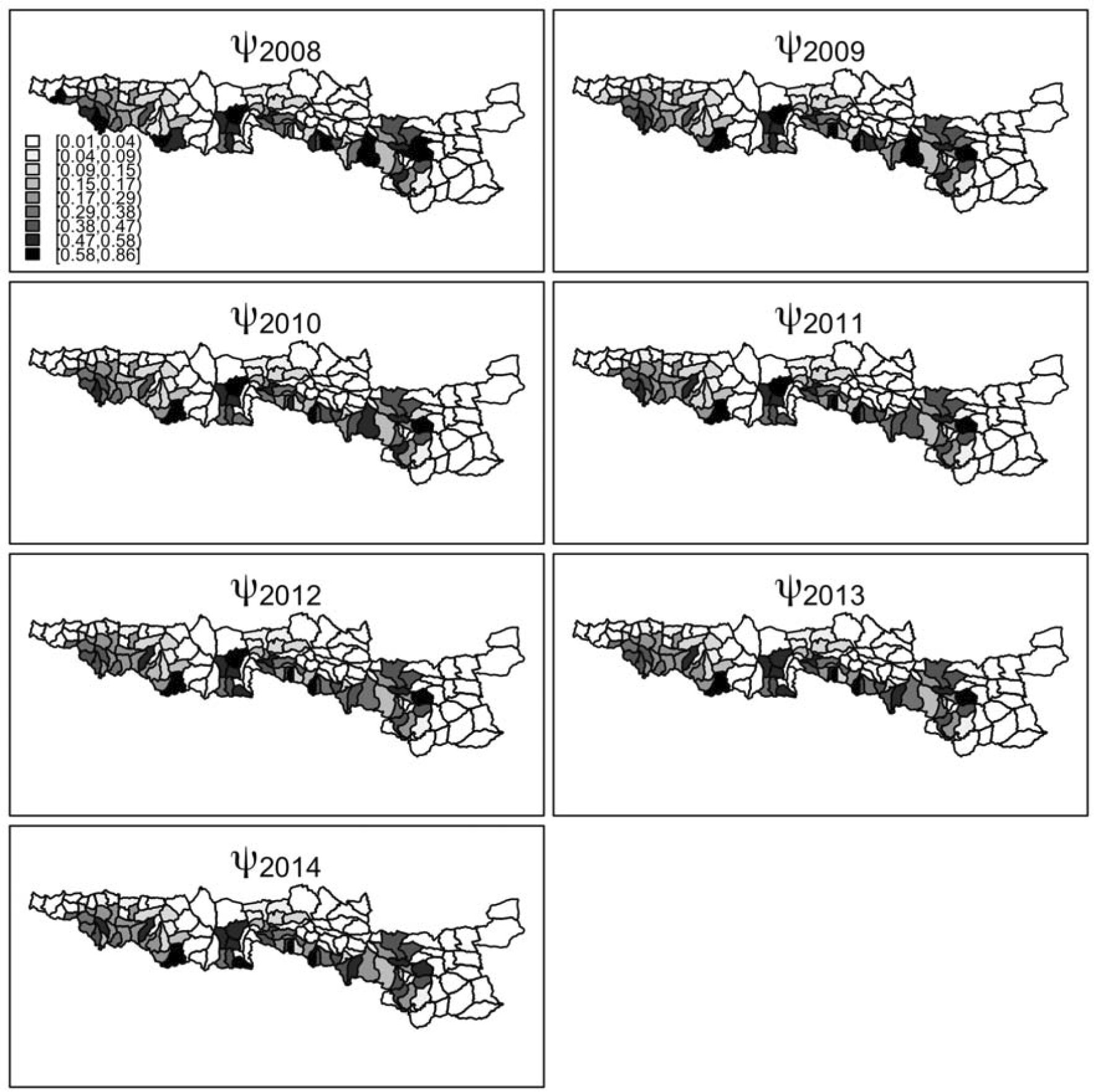
Maps of the yearly occupancy probability ψ_t_ from t = 2008 to t = 2014 in the various mountain subsections of the French Pyrenees obtained using the model-averaged parameter estimates and the formula ψ_t+1_ = (1-ψ_t_) γ + ψ_t_ (1-ε) (MacKenzie *et al.*, 2002). Covariates were set at their mean.

Occupancy in the West of the Western population core (Southwest of the Pyrénées-Atlantiques) also decreased while it remained constant in the East of that core (Southeast of the Hautes-Pyrénées). Overall, a decrease in habitat use by the bear between 2008 and 2014 was detected (p-value < 0.01 for all models in Table 3), with no new areas being colonized while others clearly went extinct.

## Discussion

### 1. Environmental and anthropogenic effects on model parameters

Human density had a strong, negative effect on initial occupancy probability ψ, with the least densely populated areas being the most likely to be used by bears. This result confirms previous analyses suggesting that bears tend to live far from the areas with the most intense human activity (Long *et al.*, 2010, Martin *et al.*, 2010). Several factors such as the habituation of the bears to human presence (Wheat and Wilmers, 2016) or the need for female bears to shield themselves from sexual conflict (Steyaert *et al.*, 2016) may mitigate this effect – but the small current size of the Pyrenean brown bear population limits the immediate relevance of these factors as bears tend to disperse further at low densities, lowering the encounter rate of other individuals and for females the risk of sexually selected infanticide (Stoen *et al.*, 2006). These results confirm that anthropogenic effects supersede natural elements when it comes to habitat selection by brown bears (Nellemann *et al.*, 2007).

Contrary to what we were expecting, human density was negatively correlated with the probability of extinction. A possible explanation is the influence of demographic stochasticity in small populations (Gabriel and Bürger, 1992) which gives more weight to extinction events. In our study, human density was lower in the Southeast of Ariège and Southwest of Aude and Pyrénées-Orientales (Figure A1) than it was in the other areas with high occupancy probability (Figure 4B), and was the place of several local extinction events in years 2010 and 2011 (Camarra *et al.*, 2012).

Finally, we found a positive correlation between the detection probability and roughness. A rougher terrain funneling pathways of bears and humans may explain this pattern. The same funneling effect might explain why signs of bears were easier to detect (when the species was present) in small subsections than in large ones. Overall, species detection was imperfect and estimated below 0.6, therefore confirming the need to correct for it to avoid underestimating occupancy.

### 2. Brown bear habitat use in the French Pyrenees

The occupancy maps for bears in the Pyrenees clearly showed the existence of two independent population cores, one located in the West and another in the Center of the Pyrenees (Figure 4B, Figure 5). The two cores remained unconnected during the timespan of the study. The dynamics of occupancy over the study period (Figure 5) showed that habitat use significantly decreased overall. In particular, the extinction of the Eastern part of the Central core is consistent with the lack of bear tracks found in Southeast Ariège and Southwest Aude and Pyrénées-Orientales (Figure 1) since 2011 (Camarra *et al.*, 2012). These results demonstrate the usefulness of dynamic occupancy models to highlight trends in habitat use that cannot be identified by static species distribution models (MacKenzie *et al.*, 2003).

The fact that we found many mountain subsections with a high occupancy probability in the Western core despite the fact that only 2 to 3 bears were estimated to live there between 2008 and 2014 (Piédallu *et al.*, 2016b) suggests a violation of the closure assumption between our secondary occasions (July-November), because there were not enough bears in the population core to occupy all subsections at the same time. This means that we estimated the habitat use by brown bears instead of the actual occupancy. For species that can attack livestock, presence does not have to be permanent to be a source of conflict, and therefore habitat use remains a relevant indicator in the case of large carnivores often characterized by their relatively large home ranges (Gittleman and Harvey, 1982) and their use of large areas without actually occupying much land at any given time.

### 3. Implications for human-wildlife conflict mitigation

We anticipate that our results will be useful as part of the “scientific evidence gathering” that is required for conflict mitigation (Redpath *et al.*, 2013). Attacks on livestock are one of the main causes of the negative attitudes towards carnivore presence in general (Kaczensky, Blazic and Gossow, 2004, Sponarski *et al.*, 2013) and towards brown bears in the Pyrenees in particular (Piédallu *et al.*, 2016a). There is an interest in mapping the areas which are more likely to host bears in the present and the future, and as such the “attack hotspots” (Miller, 2015). It could also be combined with a mapping of attitudes towards brown bears (Piédallu *et al.*, 2016a) to identify areas that combine positive attitudes towards bear presence and low attack risk, and as such could be primary targets of future management decisions. This might be the first step towards the development of socio-ecological models designed to mitigate human-wildlife conflicts (Aswani, 2011, Dupont *et al.*, 2011, Estoque and Murayama, 2014).

## Acknowledgments

We are grateful to the volunteers of the Brown Bear Network and the ONCFS Bear Team for collecting and sharing precious data and knowledge on the Pyrenean brown bears.

## Author contributions

Conceived and designed the experiments: BP, PYQ, OG. Performed the experiments: BP, NB, AG, CM, PYQ. Analyzed the data: BP, OG. Contributed reagents/materials/analysis tools: BP, PYQ, NB, AG, CM, OG. Wrote the paper: BP, PYQ, OG.

## Biographical sketches

Blaise Piédallu is a population ecologist interested in human-wildlife conflicts with a focus on large carnivores. Pierre-Yves Quenette is an ecologist who leads the ONCFS brown bear program. Nicolas Bombillon is an ecologist interested in wildlife conservation. Adrienne Gastineau is an ecologist interested in the behavior of large carnivores. Christian Miquel is a population geneticist interested in promoting non-invasive monitoring methods. Olivier Gimenez is a biostatistician interested in population dynamics of large carnivores.

## Supplementary materials

**Figure A1:**
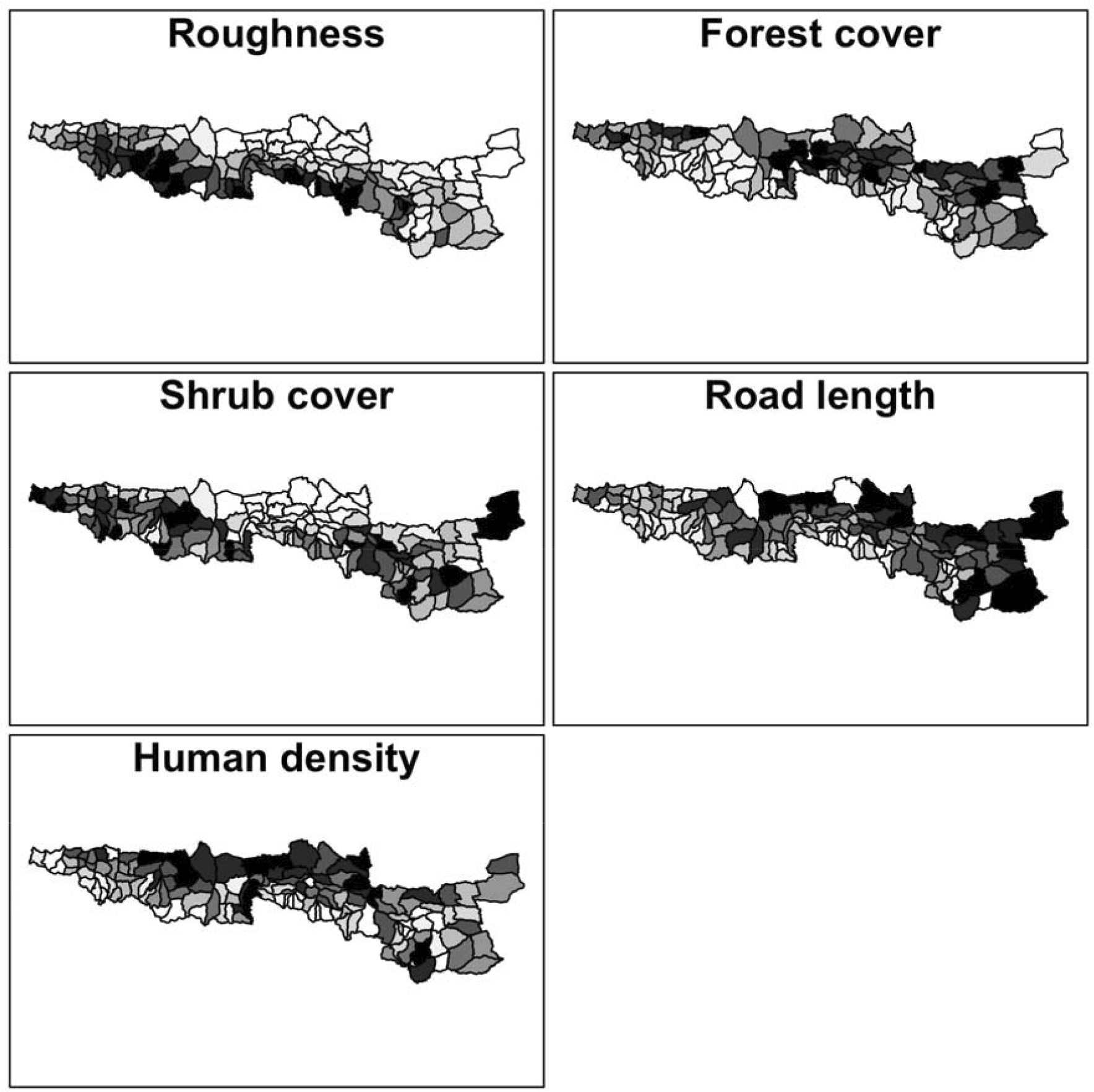
Maps of the five environmental or anthropogenic covariates in the mountain subsections of the Pyrenees that were used to build the occupancy models (see also Table 1).

**Figure A2:**
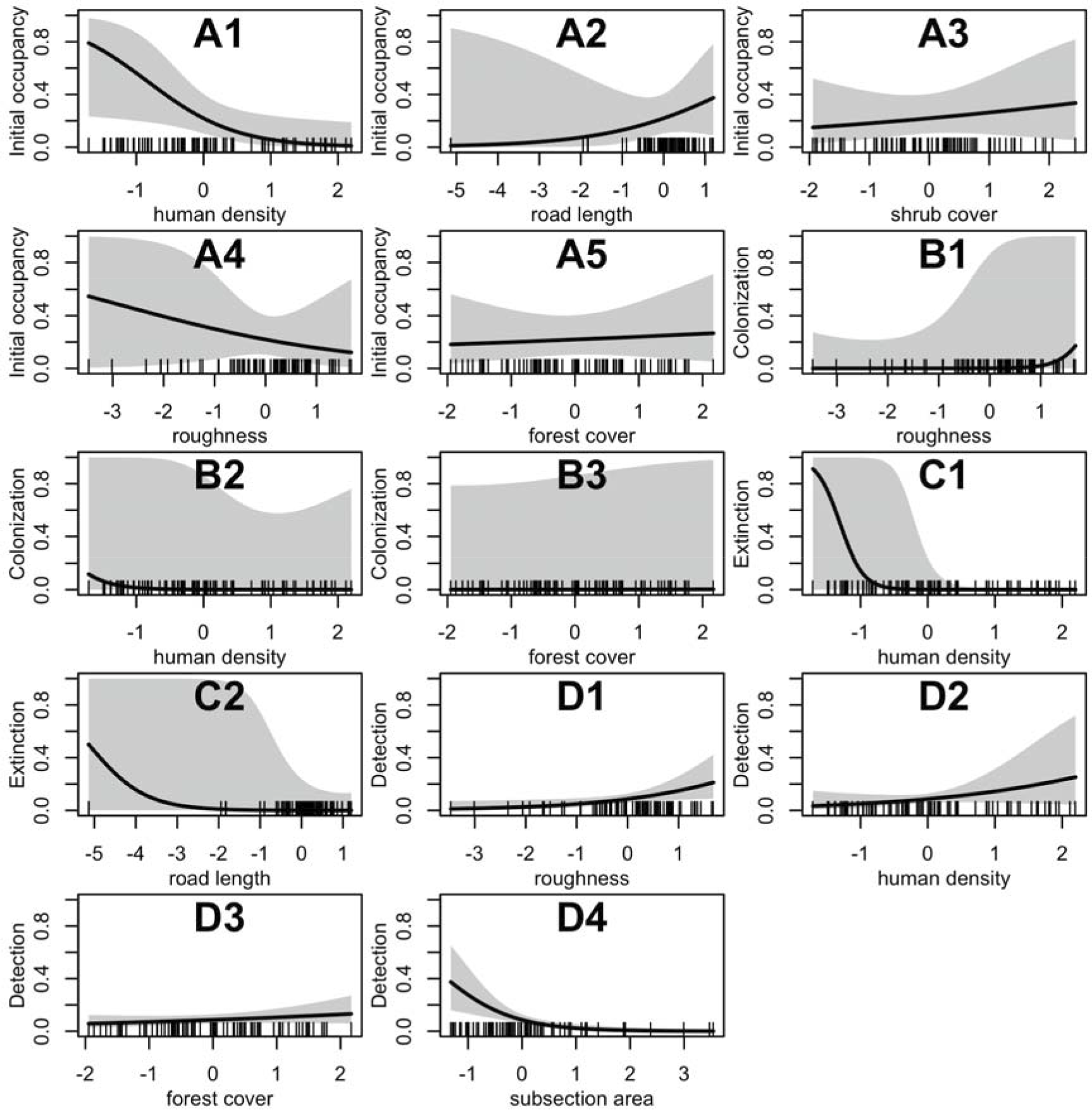
Relationships between model-averaged parameter estimates and standardized covariates. A1-A5 is for initial occupancy, B1-B3 for colonization, C1-C2 for extinction and D1-D4 for detection. For each relationship, the non-focal covariates were set at their mean. The dashes on the x-axis indicate the observed covariate values.

